# EBADIMEX: An empirical Bayes approach to detect joint differential expression and methylation and to classify samples

**DOI:** 10.1101/401232

**Authors:** Tobias Madsen, Michał Świtnicki, Malene Juul, Jakob Skou Pedersen

## Abstract

DNA methylation and gene expression are interdependent and both implicated in cancer development and progression, with many individual biomarkers discovered. A joint analysis of the two data types can potentially lead to biological insights that are not discoverable with separate analyses. To optimally leverage the joint data for identifying perturbed genes and classifying clinical cancer samples, it is important to accurately model the interactions between the two data types.

Here, we present `EBADIMEX` for jointly identifying differential expression and methylation and classifying samples. The moderated t-test widely used with empirical Bayes priors in current differential expression methods is generalised to a multivariate setting by developing: (1) a moderated Welch t-test for equality of means with unequal variances; (2) a moderated F-test for equality of variances; and (3) a multivariate test for equality of means with equal variances. This leads to parametric models with prior distributions for the parameters, which allow fast evaluation and robust analysis of small data sets.

`EBADIMEX` is demonstrated on simulated data as well as a large breast cancer (BRCA) cohort from TCGA. We show that the use of empirical Bayes priors and moderated tests works particularly well on small data sets.

## 2 Introduction

Accurate identification and robust modeling of cancer biomarkers can provide key insights to improve cancer diagnosis and treatment (Smith *et al.*, 2014). Integrated analysis of biomarkers, and the derived increase in statistical power, is believed to hold more information and enable a performance gain, compared to separate analyses of individual biomarkers (Kristensen *et al.*, 2014). Careful modeling of any dependencies between the data types is naturally important. DNA methylation and its regulation of gene expression levels has long been known to play an important part in the development and progression of cancer (Levenson, 2010). Here, we present `EBADIMEX` (<u>E</u>mpirical <u>Ba</u>yes for <u>Di</u>fferential <u>M</u>ethylation and <u>Ex</u>pression) to jointly identify differential expression and methylation, and to classify samples as either normal or tumor. We apply the method to a breast cancer (BRCA) cohort from The Cancer Genome Atlas (TCGA) (http://cancergenome.nih.gov) with available gene expression and methylation data.`EBADIMEX` is freely available as an R-package (appendix B).

DNA methylation plays a critical role in normal cell development and in shaping cell identity (Smith and Meissner, 2013). Along with histone modifications, DNA methylation is one of the main regulatory epigenetic mechanisms. The paradigm is that promoter hypermethylation silences gene expression, by directly or indirectly inhibiting transcription factor binding (Kass *et al.*, 1997; Brenet *et al.*, 2011; Jones, 2012). Conversely, gene body hypermethylation seems to correlate positively with transcriptional activity, although the causative mechanisms are poorly understood (Jjingo *et al.*, 2012; Yang *et al.*, 2014). The silencing of tumor suppressor genes through promoter hypermethylation is established as a pathway in carcinogenesis (Jones and Baylin, 2007; Esteller, 2008), and the biomarker potential of DNA methylation in cancer is reported in numerous studies (Ma *et al.*, 2016; Strand *et al.*, 2014; Zhong and Cen, 2017). Furthermore, gene body hypomethylation in a panel of genes was recently suggested as an important hallmark of cancer (Mendizabal *et al.*, 2017). Various specialized methods to detect differential methylation exist. These methods typically perform single-probe analysis or detect differentially methylated regions (DMR), where the breakpoints of the DMR regions are learned from data (Aryee *et al.*, 2014; Morris *et al.*, 2013; Wu *et al.*, 2013).

High-throughput sequencing of RNA molecules (RNA-seq) has gradually overtaken the role of microarrays in quantifying gene expression levels. Multiple methods for detecting differential expression between experimental or biological conditions in RNA-seq data exist. `DESeq2` (Love *et al.*, 2014), `edgeR` (McCarthy *et al.*, 2012), and `limma` (Smyth *et al.*, 2004; Ritchie *et al.*, 2015) are examples of such methods that all use an empirical Bayes approach to share information across genes and obtain regularized estimates of the gene-specific expression variation. By sharing information across genes, the use of empirical priors increases the power to detect differential expression, particularly when the number of samples or replicates is small. `DESeq2` and `edgeR` both model the RNA-seq read-counts directly using a negative-binomial model. `limma` was originally designed for the analysis of microarray data, which, in contrast to RNA-seq data, gives a continuous measure of expression for each gene. By log-transforming normalized RNA-seq data, `limma` has subsequently been adapted to accommodate RNA-seq data.

Methylation and expression data can be integrated by combining p-values from methods specialized for analyzing the two data types separately, e.g. by using Fisher’s method (Fisher, 1932). However, this approach rests on the typically invalid assumption of independence. Another integration approach is to use machine learning methods such as random forest or regularized regression methods, while not explicitly accounting for dependencies between data types (List *et al.*, 2014). `PINCAGE` (świtnicki *et al.*, 2016) uses a probabilistic graphical model to explicitly model the interaction between methylation and expression measurements. With `PINCAGE`, the gene-specific distributions are modelled and learned using non-parametric kernel-based methods. Probabilistic models provide a high degree of flexibility and are highly interpretable. However, kernel density estimation in high dimensions suffers from the curse-of-dimensionality. Using an example with Gaussian kernels, it can be shown that the required number of samples to obtain a fixed level of precision increases exponentially with the number of dimensions (Scott, 2008, chapter 7).

Here, we propose a parametric probabilistic model to integrate expression and methylation data. We use empirical Bayes to obtain regularized variance and covariance estimates, generalizing the approach used by `limma` to multiple dimensions. Specifically, we develop (1) a moderated Welch t-test for equality of means with unequal variances; (2) a moderated test for equality of variances; and (3) a multivariate test for equality of means with equal covariance matrices. Cancer genomics data often display large amounts of heterogeneity, and to achieve robustness to outliers, methods from robust statistics, particularly winsorization and the Huber loss, are employed (section 6.1 and appendix A.1).

To illustrate the performance of `EBADIMEX`, we classify tumor and adjacent normal breast cancer samples from TCGA. We increase the difficulty of the classification problem by using fewer genes and fewer samples to train the model. `EBADIMEX` consistently performs better in classifying samples and identifying genes with differential methylation and expression than similar methods, which neither share information across samples using empirical Bayes nor correct for outliers. We observe this for all subsample sizes, but the difference is more pronounced for small subsamples.

`EBADIMEX` is not restricted to work with RNA-seq and methylation data, but can be extended to other data types, so long as the same measurements are performed for each of a moderately large number of units. Empirical Bayes posits an informative prior which is estimated from data, and the uncertainty on the location of this prior should be minimal compared to the spread of the prior. This is typically the case if the prior is based on a couple of hundred units, although it depends on the exact setting.

In the following sections we first discuss the input data for `EBADIMEX` and its necessary preprocessing, along with methods to increase robustness to outliers. Next, the model behind `EBADIMEX` and its underlying assumptions are presented. The mathematical formulations of hypotheses regarding differential expression and methylation are stated along with the details of statistical tests. An overview of the full analysis pipeline is given, followed by descriptions of methods to validate the performance of `EBADIMEX`.

## 3 Input data for EBADIMEX

The input data for `EBADIMEX` is expression data and DNA methylation measurements. The DNA methylation data should contain measures of the average methylation at genomic CpG sites in the promoter and the gene body of a gene. Such measurements can be obtained with bisulfite sequencing or array-based techniques. The use of `EBADIMEX` is illustrated with data from the widely used Illumina Infinium 450K methylation array (Bibikova *et al.*, 2011). For example, TCGA has collected methylation profiles of almost 9, 000 samples using the array (Grossman *et al.*, 2016). Here, we analyze the publicly available TCGA breast cancer samples, where both RNA-seq expression data and DNA methylation measurements are available. The data set consists of 84 normal samples and 781 tumor samples (Weinstein *et al.*, 2013).

In RNA-seq experiments, a small fraction of the genes often contribute a large fraction of the reads. If the data is normalized by the total read count, differential expression of one of these genes will affect all other genes. To alleviate this problem, we use upper-quartile normalization. We further log-transform these normalized values to reduce the skewness, that such count data would otherwise display. After these pre-processing steps, we denote the expression level of gene *g* in class *c* and sample *i* by *E*_*gci*_.

For each gene there is a number of associated methylation probes. We use the annotations of the probes provided by the Illumina development team to define promoter and genebody associated probes for each gene (Bibikova *et al.*, 2011). In the promoter definition we include the region from 0-1500 bp upstream of the transcription start site (TSS), 5’ UTR and 1st exon. The 1st exon is included in the promoter definition, as it has been shown that methylation of the 1st exon has similar regulatory consequences as promoter methylation (Brenet *et al.*, 2011). The gene body is defined as the remaining transcribed region (Figure 1A-B). We calculate the average promoter and gene body M-value, defined as the log_2_ ratio between the number of methylated and unmethylated probes, potentially adding a pseudo count *γ*,

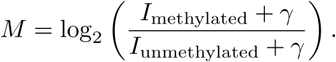

**Figure 1:**
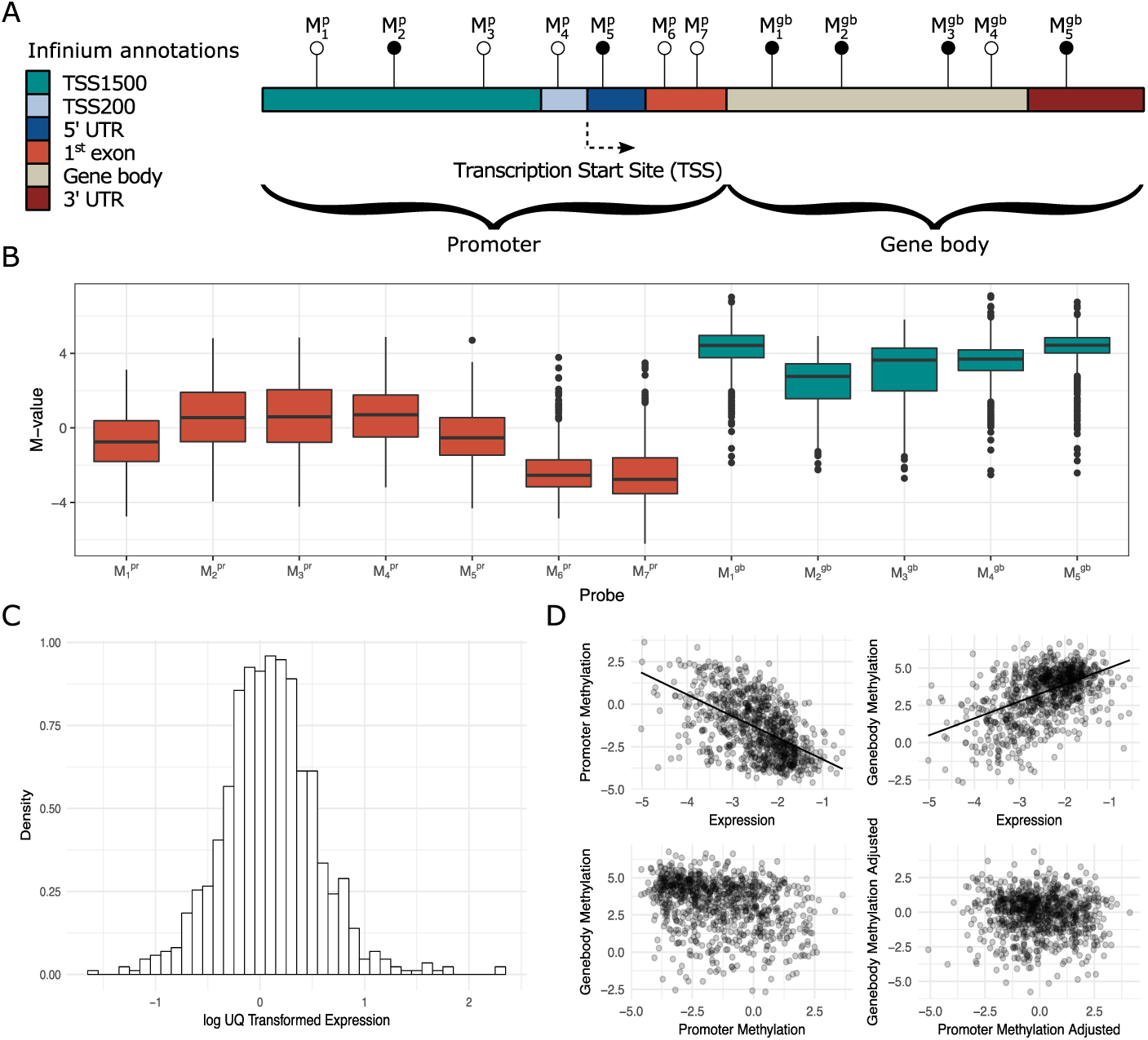
Data overview. **(A)** Methylation probes are classified as belonging to promoters and gene bodies, as defined by Illumina. The 1,500 bp upstream of the transcription start site (TSS), the 5’ UTR and the 1st exon are included in the promoter region definition. The remaining coding region and the 3’ UTR are included in the gene body region definition. **(B)** Probe measurements for a single gene across the BRCA cohort. We average over the promoter (red) and gene body probes (green), respectively. **(C)** Distribution of expression levels for a single gene after upper-quartile normalization and log transformation. **(D)** Correlation between expression and promoter- and gene body methylation. Generally, promoter methylation and expression are negatively correlated, while gene body methylation and expression are positively correlated. Also shown is promoter methylation against gene body methylation, both raw and adjusted for the expression level.

M-values are typically preferred over the raw fraction of methylated probes because of the generally heteroscedastic nature of the raw fractions (Du *et al.*, 2010).

We denote the average promoter and gene body methylation of gene *g* in class *c* and sample *i* by 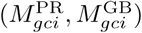.

### 3.1 Data transformations

To cater for the large amount of heterogeneity observed in the expression patterns of cancers, we take additional measures to ensure robustness, as outlined below.

#### Pre-filtering

Only genes that are consistently expressed across samples are included in the analysis. It is a common practice used both for RNA-seq and microarrays to filter out genes that are very lowly expressed. For RNA-seq, low expression levels manifest themselves as many samples with zero read-count. First, such genes are likely not implicated in differentiating cancers, and second, when log-transforming small read counts they are not well approximated by a normal distribution and thus violate the distributional assumptions of linear normal models used later.

#### Normalization

Normalizing RNA-seq data has been identified as a crucially important step in differential expression studies (Bullard *et al.*, 2010). Some of these procedures address the problem of controlling for transcript length in deciding the actual relative abundance between two transcripts, leading to the widely used FPKM measures. This problem is irrelevant for `EBADIMEX`, as it only handles the relative difference between samples for individual transcripts.

However, another important problem is that a small group of genes make up the bulk of the RNA-seq reads (in the TCGA breast cancer cohort, 6.6% of the genes contribute half the reads). If one or more of these genes are differentially expressed, normalizing by total read count will dilute the differential expression signal. Furthermore, false positive differential expression can be introduced for more lowly expressed genes. Here, we use upper-quartile normalization to avoid the adverse effects of total count normalization (Figure 1C).

#### Winsorization

A final important step is to ensure robustness to outliers, which are frequent in cancer genomics. Where outliers do not pose much risk for kernel-based methods, parametric methods are generally sensitive to outliers. We apply winsorization, which is an iterative procedure that cap a data vector such that no value is more than a pre-specified number of standard deviations away from the mean (Huber and Ronchetti, 2009).

An important feature of the data is that expression and methylation correlate: generally expression correlates negatively with promoter methylation and positively with gene body methylation. However, we observe large intergene differences. Moreover, gene body-and promoter methylation are typically slightly positively correlated after adjusting for the effect of expression. (Figure 1D)

## 4 Statistical models of EBADIMEX

At a high level we model first the marginal distribution of the expression data and then the joint distribution of the gene body and promoter methylation measurements conditional on the expression data.

### 4.1 Expression model

The expression model of `EBADIMEX` is similar to the model used in the `limma` software package (Smyth *et al.*, 2004; Ritchie *et al.*, 2015), where a linear normal model is fitted for each gene, but share information across genes using empirical Bayes. A prior for variance is learned across all genes. This increases the statistical power to detect genes with large effect sizes but high estimated variances, at the expense of genes with low effect sizes and small estimated variances. In the present work we generalise the previously developed moderated t-test (Smyth *et al.*, 2004), used with empirical Bayes priors in expression analysis (Smyth *et al.*, 2004; Ritchie *et al.*, 2015): We thus develop a moderated Welch t-test to test the hypothesis of equal means with unequal variances and a moderated F-test to test the hypothesis of equal variance.

The model introduced in `limma` assumes the following distribution of the log-expression level, *E*_*gci*_, of gene *g* in class *c* for sample *i*:

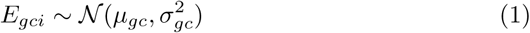

Where

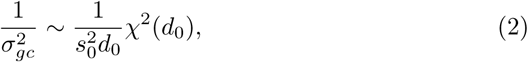

i.e. 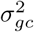 has an inverse-*χ*^2^ distribution.

Realistic models of the underlying count data are heteroscedastic, and this feature carries over to the log-transformed data. As an example, using the delta method, it can be seen that the log-transform of a Poisson distributed variable has a variance that is inversely proportional to its mean. Consequently, the hyperparameters *d*_0_ and *s*_0_ are functions of the expression level. To estimate these functions we first bin the genes into groups with similar expression levels. Next, we use an empirical Bayes approach to learn a pair of hyperparameters *d*_0_ and 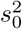, for each group. Finally we interpolate the parameters of for each expression level (Figure 2A, section 6.6).

**Figure 2:**
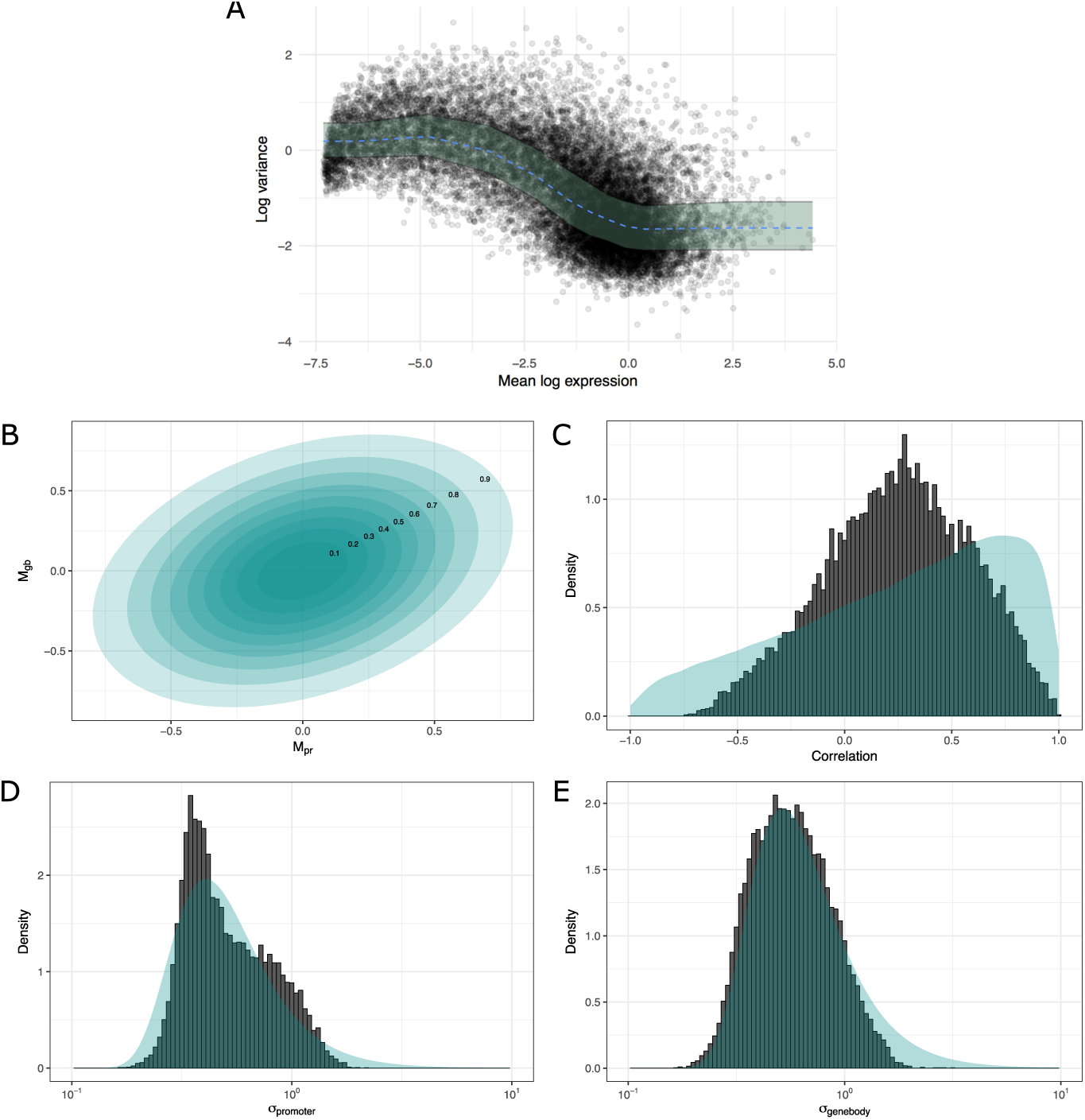
Priors. **(A)** The prior variance for expression depends on the expression level. Using empirical Bayes we estimate the variance prior. The empirical variance estimates are plotted (black dots) along with the median of variance prior (dotted blue line). The shaded region represents the first and third quartile of the variance prior. **(B)** The mean covariance matrix for the inverse-Wishart distribution used as prior for methylation. **(C-E)** The prior distribution of correlation, standard deviation for promoter methylation and standard deviation for gene body methylation.

We are interested in not only the hypothesis of shift in location, *H*_*µ*_: *µ*_*g*1_ =*µ*_*g*2_, but also the hypothesis of variance homogeneity, 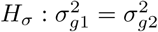. Moderated versions of the classical t-test and novel moderated versions of the Welch t-test and the F-test of variance homogeneity are used to test the two hypotheses (sections 6.2, 6.3 and 6.4). In the moderated tests, a regularized version of the variance estimate replaces the regular estimate. The advantage of these tests is that they increase power, but also that they emphasize effect size over small variance.

### 4.2 Methylation model

For a particular gene we model the two methylation values (gene body and promoter) as a bivariate normal distribution, where the mean depends linearly on the expression level and on the group (Figure 2A).

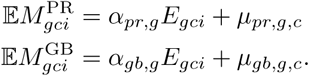

Typically, the measurements from the two probe types are positively correlated, but due to a large variation between genes, some genes display a negative correlation as well (Figure 2C). The correlation between probe types is determined by the covariance matrix, Σ, of the bivariate normal distribution (Figure 2B-E). The inverse wishart distribution has four free parameters, of which three capture the mean of the two variances and the correlation, and one captures the peakedness of the distribution (i.e. how informative the prior is). Therefore, it is slightly inflexible when it comes to modeling the correlation accurately. The prior is conservative with the amount of information we have on the distribution of the correlation coefficient (Figure 2C).

Collecting all parameters specifying the mean value structure in a single matrix,

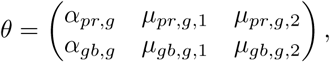

the model can be written as

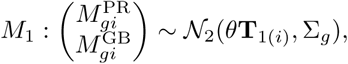

where 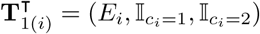 is the *i*’th row of the N by 3 design matrix **T**_1_,N being the total number of samples.

The covariance matrix, Σ, is assumed to be distributed according to an inverse-Wishart distribution

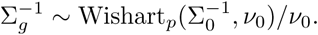

This is analogous to the use of the inverse-*χ*^2^ distribution for the variance parameter in the expression model (equation (2)). As before, the parameters Σ_0_ and *v*_0_ are learned using empirical Bayes (section 6.7).

The relevant hypothesis for the methylation model is whether the two groups have the same mean after the controlling for expression:

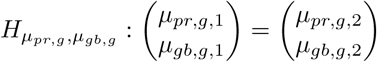

As we are now testing for equality of two parameters simultaneously a moderated version of the F-test is used to test the hypothesis (section 6.5).

## 5 EBADIMEX pipeline

The analysis pipeline is summarized in this section, and a schematic overview of the steps in `EBADIMEX` is given in Figure 3.

**Figure 3:**
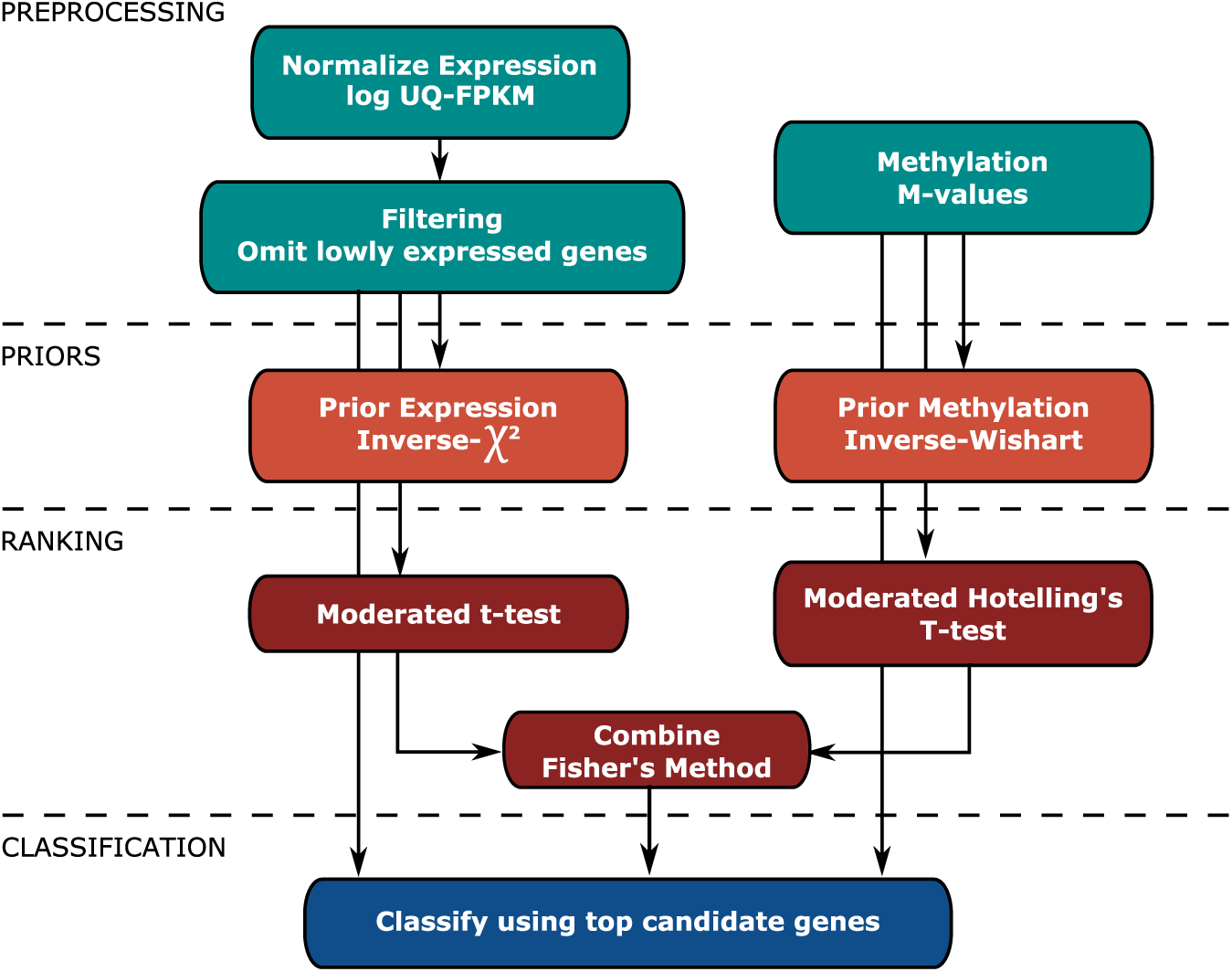
Schematic overview of the analysis pipeline.

### Preprocessing

We perform a number of data processing steps: If the RNA-seq data is in the form of raw read counts for each gene, we normalize the data. The options for normalization in the `EBADIMEX` R-package include total count-and upper quartile normalization. By default, upper quartile normalization is used (appendix A.2). We proceed with filtering out genes with ubiquitously low expression. This is done by dichotomizing based on median expression level of the individual genes. The threshold is guided by fitting a two-component normal mixture to the median expression levels; genes whose median expression level falls in the lower of the two components are disregarded in subsequent analysis (appendix A.3).

### Priors

After filtering the genes, raw estimates of the expression variance and the methylation covariance matrices are obtained for each gene. Based on the distribution of these we learn the empirical prior distributions (sections 6.6 and 6.7).

### Ranking

Next, we use moderated test statistics to determine differential regulation. Genes are ranked according to their p-value. Alternatively, as an option in the `EBADIMEX` R-package, the data can be fit using regularized estimates after which we are able to calculate the Kullback-Leibler divergence between the fitted distributions (appendix A.4).

### Classification

Finally, we classify samples using a likelihood criterion:

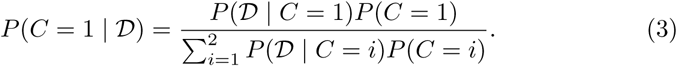

where 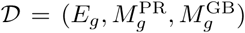 is the data for a single gene, *g*. The classifiers are evaluated using the area under the ROC curve (AUC) in an n-fold cross validation scheme. The AUC performance measure is agnostic to the actual class probabilities, but only considers the ordering of samples. The Bayes Factor (BF) is defined as

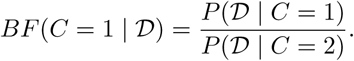

As the BF is a monotone function of *P* (*C* = 1 | 𝒟), given in equation (3), it leaves the measure of AUC invariant. Furthermore, using the BF there is no need to specify the prior class probabilities *P* (*C* = *i*). We therefore use the BF as predictor of the class label. The Bayes factor can be computed as

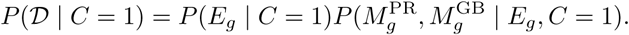

Here, the first term is

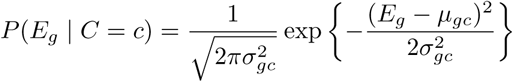

and the second term is

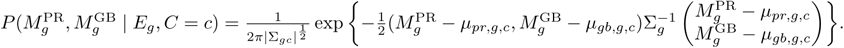

For both terms, regularized variance and covariance estimates are used.

## 6 Statistical details

### 6.1 Winsorization

Let *x* = (*x*_1_, *…, x*_*N*_) be a data vector. Winsorization is a procedure that caps the original data for small and large values:

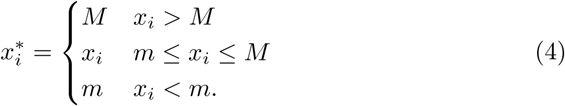

There are different approaches to determine *m* and *M*. One approach is setting *m* and *M* at the 5th and 95th percentile, respectively, thus ensuring that 10% of data is capped. Here we use an iterative procedure described in Huber and Ronchetti (2009):

1. Set *m* = *-−* and *M* = *−*.

2. Compute *x*^*\**^ using (4).

3. Compute the empirical mean, *µ*_*x*_*\**, and the standard deviation, *Σ*_*x*_*\**, of *x*^*\**^.

4. Set *m* = *µ*_*x*_*\* − k * σ*_*x*_*\** and *M* = *µ*_*x*_*\** + *k * σ*_*x*_*\**.

5. Repeat from step 2 until convergence.

The value *k* determines how aggressively the data is capped. For our analysis *k* = 2.5 is used.

Estimates based on winsorized data has a smaller effective degree of freedom than estimates based on the raw data. We need to correct for this in estimates and tests. The idea proposed by Dixon and Tukey (1968) is to scale the t-test statistic by a factor 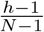, where *h* is the number of observations left unaltered by winsorization. In the subsequent t-test, the degrees of freedom is similarly adjusted to *h* – 1 rather than *N* – 1. The effect of scaling the t-test statistic can also be obtained by scaling the estimate of the standard deviation (Huber and Ronchetti, 2009).

### 6.2 Moderated t-test for equality of means

Let *X*_*ij*_ *∼ 𝒩* (*µ*_*i*_, *σ*^2^), *i* = 1, 2, *j* = 1, *… n*_*i*_ be two independent normal samples with common variance. Furthermore, let *σ*^2^ follow a scaled inverse-*χ*^2^ distribution,

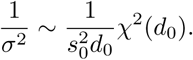

We want to test the hypothesis *H*_*µ*_: *µ*_1_ = *µ*_2_ of common mean. The means are estimated by

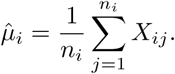

These estimates are independent of the unbiased variance estimate given by

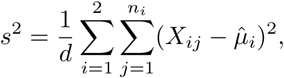

where *d* = *n*_1_ + *n*_2_ – 2. The moderated t-test uses a regularized estimate of *σ*^2^,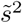:

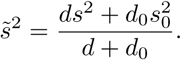

Replacing *s*^2^ with 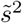 in Student’s t-test we obtain the moderated t-test statistic,

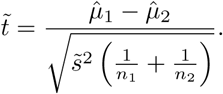

Under the hypothesis 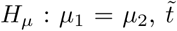 is t distributed with *d* + *d*_0_ degrees as is shown in Smyth *et al.* (2004) using transformation of variables and density functions. Equivalently, 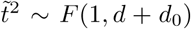. Here we provide a simpler proof. First, note that

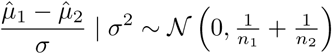

and is conditionally independent of

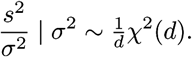

Since the conditional distributions do not depend on *σ*^2^, the stated distributions apply unconditionally. At the same time,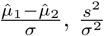, and *σ*^2^ are all independent. Next we see that,

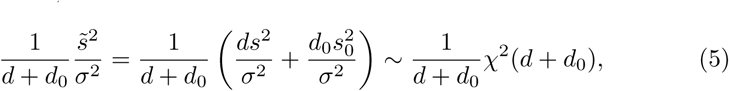

since the term in the parenthesis is the sum of two independent *χ*^2^ distributions with *d* and *d*_0_ degrees of freedom, respectively. Finally, writing

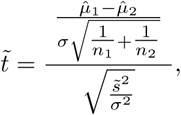

we see that 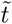 is the ratio between a standard normal distribution and the square root of an independent 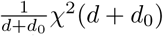, which characterizes the *t*-distribution with *d* + *d*_0_ degrees of freedom.

### 6.3 Moderated Welch t-test for equality of means with unequal variances

As an alternative to the moderated t-test, we also consider a moderated Welch t-test for the case of unequal variance between two groups. Demissie *et al.* (2008) propose the following test:

Consider *X*_*ij*_, *i* = 1, 2 and *j* = 1, *…, n*_*i*_ and assume

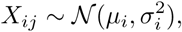

Where

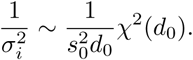

The usual estimators of the variances are denoted by

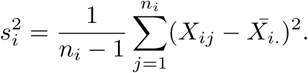

The moderated Welch t-statistic is

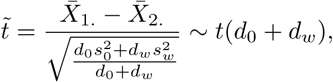

where

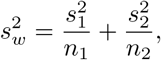

and

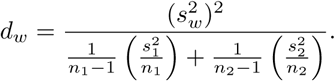

The moderated Welch t-test is not well founded mathematically, and, indeed, our simulations show that it might be poorly fit by the t distribution (Supplementary Figure S1A). We therefore propose another version of a moderated Welch t-test, where variance estimates are substituted with moderated variance estimates:

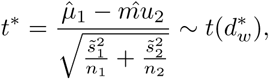

Where

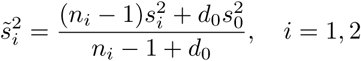

and

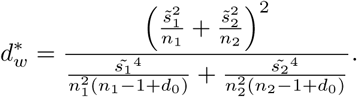

The fact that *t*\* is approximately t distributed follows from the Welch-Satterthwaite approximation. First, note that writing

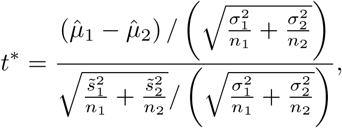

the numerator follows a standard normal distribution and is independent of the denominator. Using the Welch-Satterthwaite, the denominator is approximately 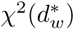 distributed, and, hence, *t*\* is approximatly t distributed with 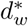 degrees of freedom. Our simulations show that this moderated Welch t-test statistic is better approximated by the t distribution (Supplementary Figure S1B). Another thing to note is that the two sample variances need not have the same prior distribution.

### 6.4 Moderated F-test for univariate equality of variances

Let 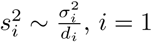, 2 and let

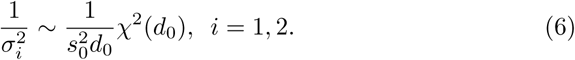

We want to test the hypothesis *H*_*σ*_: *σ*_1_ = *σ*_2_, of common variance, where the variance is drawn from the distribution in equation (6). Thus, 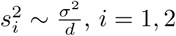 where

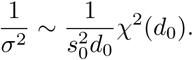

Assume, without loss of generality, *d*_1_ ≤ *d*_2_. In the moderated F statistic we use a regularized variance estimate for the smaller of the two samples,

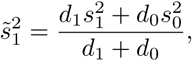

and consider the following test statistic

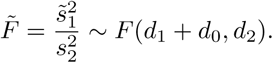

To see that 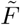 indeed follows the posited F distribution, note that with an analogous argument to that used in section 6.2, it can be shown that 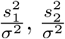 and *σ*^2^ are all independent. Using equation (5) we see that 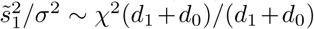 is independent of 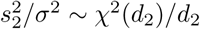, hence

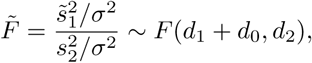

as the F distribution is characterized by the ratio of two scaled *χ*^2^ distributions.

Using moderated variance estimates for both *σ*_1_ and *σ*_2_ will not yield a proper F distribution. Regularizing either of the two variance estimates, will lead to a proper F distribution. The choice of regularizing the variance estimate for the smaller class instead of the larger class, will however give slightly more power (Supplementary Figure S2).

### 6.5 Moderated F-test for multivariate equality of means

Collect all methylation measurements in an *N ×* 2 matrix:

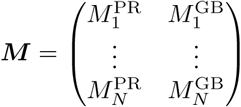

The hypothesis of no group effect in the methylation part of the model, i.e.

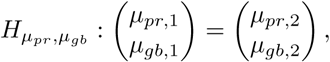

can be tested with an F-test.

First, note that the test can be phrased as a reduction from two nested multiple regression models. The design matrix for the model with group effect has rows

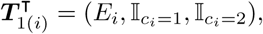

and the design matrix for the model with no group has rows

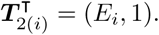

The projections onto the subspaces spanned by ***T***_*j*_ are

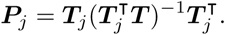

Define the SSD matrices as

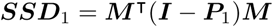

and

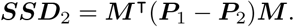

The SSD matrices are distributed according to a Wishart distribution conditional on Σ,

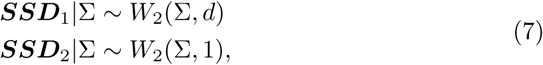

where *d* = *N* – 3.

Thinking of Σ as fixed with no prior information, the likelihood ratio statistic is

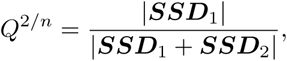

which follows the beta distribution 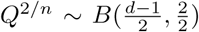. The beta distributed statistic can be transformed into an F distributed statistic:

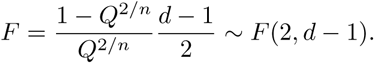

We now assume

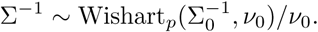

The moderated t-test can be generalized to a moderated F-test. Define first

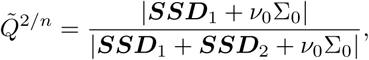

and then define the moderated F statistic,

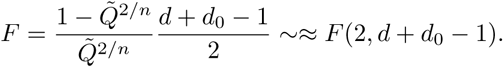

The moderated F statistic follows approximately an F distribution. It is thus computationally inexpensive to evaluate the statistical significance. The moderated F statistic is well approximated by the F distribution, as shown by simulation and by permuting real data (Supplementary Figure S3).

### 6.6 Prior Expression

The variance is assumed to follow an inverse-*χ*^2^ distribution,

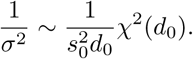

The empirical variance 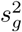 for a gene, *g*, is *χ*^2^ distributed conditional on *σ*^2^,

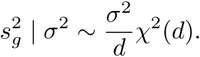

Again, 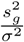is *χ*^2^(*d*) distributed and independent of *σ*^2^. We can write 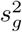 as a fraction between two independent (scaled) *χ*^2^ distributions, and it follows that 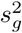 has a scaled F distribution,

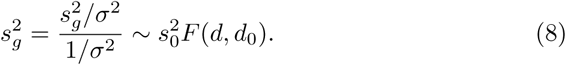

Using equation (8), the two hyperparameters,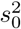 and *d*_0_, can be estimated using the method described in Smyth *et al.* (2004). The formulas are restated here for completeness.

Define 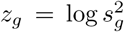. From equation (8) we see that *z*_*g*_ follows Fisher’s z distribution plus 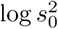. We now use moment matching to estimate *d*_0_ and 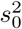. The moments of Fisher’s z distribution are

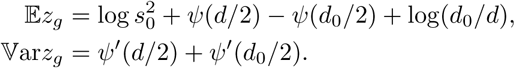

From these we see,

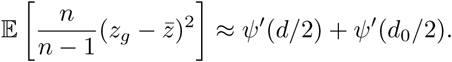

We can therefore estimate *d*_0_ by solving the following equation numerically:

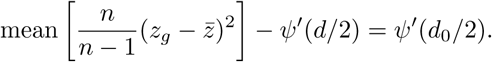

Having an estimate of *d*_0_ we solve

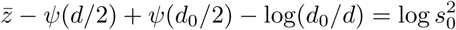

to find 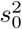.

It has been observed that the variance is dependent on the expression level, with higher expressed genes showing less variance (Love *et al.* (2014); Ritchie *et al.* (2015); McCarthy *et al.* (2012)). We therefore bin genes based on average expression level and learn the hyperparameters for each bin separately. For the analysis of a single gene we use interpolation to decide the hyperparameters (Figure 2).

### 6.7 Prior Methylation

The covariance matrix for methylation is assumed to follow an inverse-Wishart distribution,

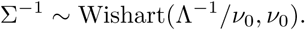

The diagonal entries in Σ then follow an inverse-*χ*^2^ distribution,

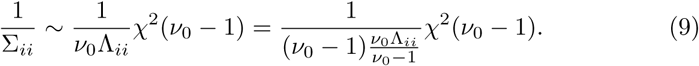

The empirical covariance matrix *S*_*g*_ for a gene, *g*, is Wishart distributed conditional on Σ,

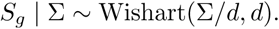

The marginal distribution of the diagonal entries is

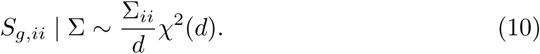

Combining equation (9) and (10), it follows using the argument presented in section 6.6 that

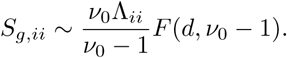

Thus, *v*_0_ and Λ_11_ and Λ_22_ can be estimated using the technique presented in section 6.6. Λ_12_ = Λ_21_ is estimated using a moment matching procedure.

## 7 Performance validation of EBADIMEX

In the following sections, methods for determining the performance of `EBADIMEX` are described. These include simulation and permutation studies as well as classification performance testing.

### 7.1 Simulation analysis

As a first mean to evaluate the performance of a test, we simulate data from the null distribution. We simulate the genes in such a way, that they have the same distribution of mean expressions as observed in the BRCA cohort. For each gene, we simulate 10 samples of which 3 are labelled as normals, and the remaining 7 are labelled as tumors. The moderated t-test is exact and the simulation thus serves as a sanity check. On the other hand, the moderated F-test is approximate. We expect that the QQ-plot for the moderated t-test displays uniform p-values. Furthermore, if the QQ-plot for the moderated F-test displays uniform p-values, it confirms that the test is applicable.

### 7.2 Permutation analysis

To evaluate if our proposed tests are robust when applied to real data, we perform permutation analyses. For each gene we randomly select 10 samples among the the tumor samples and label 3 of these as normals. The remaining 7 are labelled as tumors. The effect of using moderated tests diminish in large cohorts, and we thus only use 10 samples at a time. As the labelling is random we expect no signal of differential expression or differential methylation, and as such the p-values should be uniform.

### 7.3 Classification performance

To evaluate classification performance the following scheme is employed. First, 500 genes are selected at random, and only these genes are used for subsequent classification of samples. Next, the normal samples are divided evenly at random into *k* folds, and these folds are paired with groups of tumor samples to form *k* training datasets. To show the performance on small, but still unbalanced, data sets, only one fourth of the tumor samples are included in the training datasets. As an example, for *k* = 32, the smallest fraction of data used for training, the training data set consists of 2 – 3 normal samples and 6 – 7 tumor samples. For each of the *k* training data sets, a classification pipeline is used to classify the remaining test samples and computing the area under the curve (AUC) of the classifier. AUCs are then averaged over the *k* folds. The entire process of randomly sampling the genes used for classification and the training sets is repeated 200 times.

More than one classifier can be used on each training data for their subsequent performance comparison. For example, we compare the use of moderated and unmoderated tests. To test if a classification pipeline is significantly better than another, the Wilcoxon signed-rank test for paired samples is used.

## 8 Results

The performance of `EBADIMEX` was tested on a breast cancer cohort from TCGA. Samples were included on the basis of availability of both RNA-seq and 450K array methylation data. Following this criterion, 781 cancer samples and 84 adjacent normal samples were included.

Parameters were fit to each gene for both expression and methylation given the expression. Figure 4A illustrates the fitted distribution of expression for tumors and normals for the a highly differentially expressed gene, MATN2, and Figure 4B illustrates the fitted joint methylation distribution for the top differentially methylated gene, PKLR. Although we generally see that the empirical Bayes regularization shrinks the variance estimates in these cases, it is not immediately clear from these figures. This is explained by the relatively large amount of data used here.

**Figure 4:**
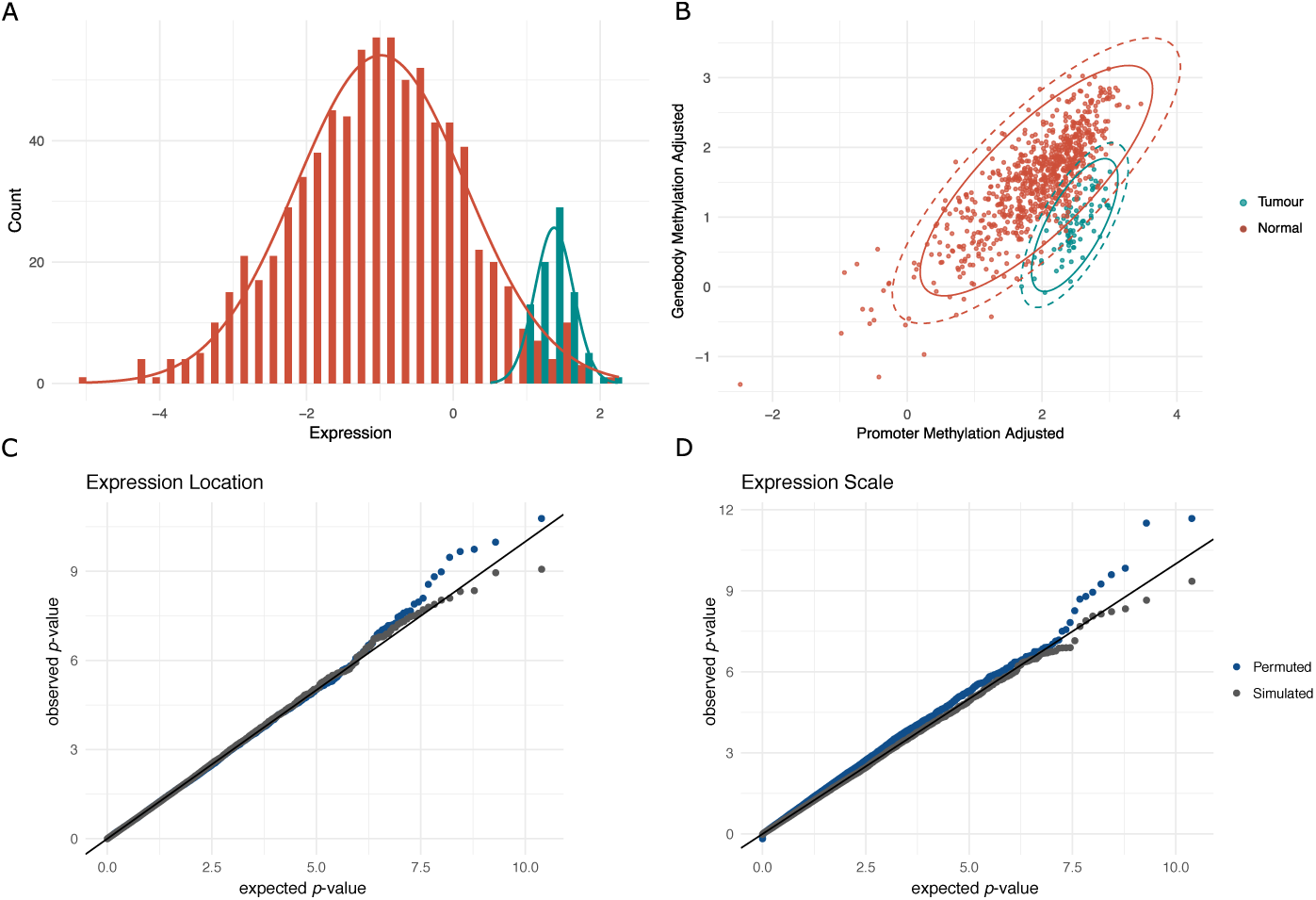
Model fit. **(A)** The distribution of expression for tumors (green) and normals (red) for the top differentially expressed gene, MATN2, in the BRCA cohort. **(B)** The joint distribution of methylation for promoter and gene body after adjusting for the effect of expression. The moderated (solid line) and unmoderated (dashed line) contour ellipses are shown. Here shown for the top differentially methylated gene, PKLR. **(C,D)** QQ-plot for the tests with permuted labels (blue) and simulated data (gray). We show the plots for the test of differential expression location and differential expression scale. QQ-plots for the remaining tests are shown in Supplementary Figures S1, S3, and S4.

To test whether the obtained test statistics were well approximated by their posited distributions, two analyses were performed: In the first analysis, data was simulated from the null model with the same number of normals and tumors in each group, and in the second analysis, 84 of the 781 tumor samples were randomly labeled as normals (sections 7.1 and 7.2). In both types of analyses, we tested for differential expression and differential methylation. Differential expression was tested with the moderated t-test for shift in location and the moderated F-test for variance homogeneity. The qq-plots obtained from these analyses show no inflation or deflation, indicating that the test statistics are well approximated by the posited distributions (Figure 4C-D and Supplementary Figure S4). Differential methylation was tested with the moderated F-test. Again, the p-values were uniformly distributed in both analyses (Supplementary Figure S3).

After the detection of differentially expressed genes, we were interested in assessing the accuracy of classification with top-ranked genes (section 7.3). Tumors and normals in the BRCA cohort can be separated almost perfectly using the top-ranked genes (Figure 4A-B and Figure 5A). To make the problem harder, the data was subsampled into successively smaller subsets. These subsets were used to detect differentially regulated genes, and the classification performance was evaluated on the remaining data. For comparison, prioritization and classification was performed using two approaches: First, with winsorization, moderated estimates and moderated tests (moderated classification). Second, without using any of the aforementioned techniques (raw classification). We sampled a subset of 500 genes and subsets of the samples of varying sizes. In the smallest subset, only 1*/*32’th of the normals, corresponding to 2 – 3 normal samples, were used. Roughly twice as many cancer samples were used. Using these samples, the genes were prioritized and the parameters of the model were learned. Then, the top-ranked gene was used to classify the remaining samples. We repeated this process 50 times. When half of the normals were used, the average AUC was 0.977 with both classification pipelines. When only 1*/*32’th of the normals were used, the average AUC with the moderated classification pipeline was 0.913, and 0.877 with the raw classification pipeline (Mann-Whitney rank sum test: *p* = 4.12 · 10^−10^) (Figure 5A). The described analysis shows that the use of robust statistics and empirical Bayes is particularly powerful for small cohorts.

**Figure 5:**
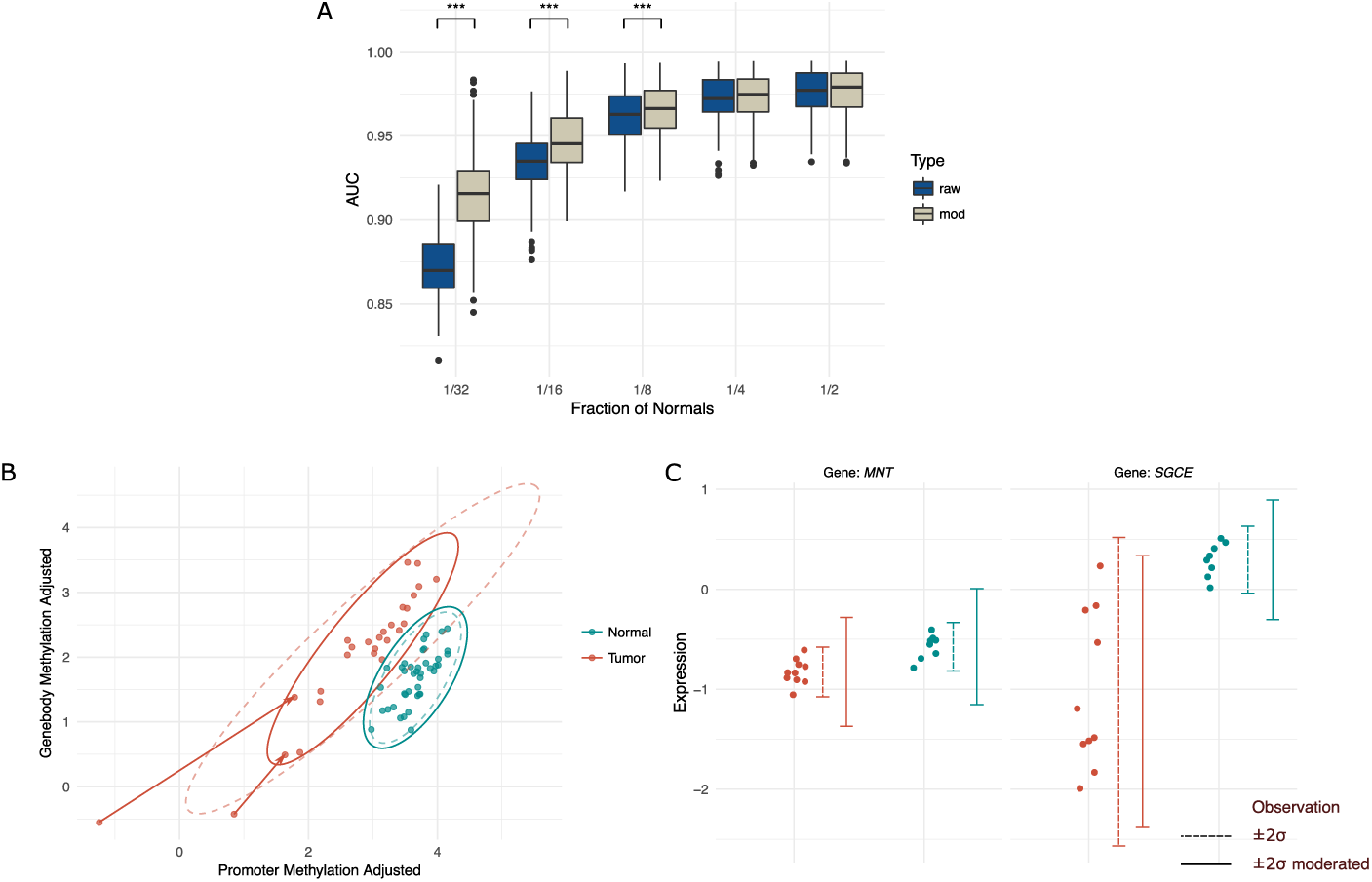
Moderated tests and classification performance. **(A)** Classification in the BRCA cohort using the top-ranked gene with respect to the p-values reported from the moderated (beige) and unmoderated (blue) tests. Likewise, the estimates used to classify the samples are either moderated or un-moderated. We use a subsample of 500 genes and progressively smaller training data sets, with the smallest (1/32th) using only 2-3 normal samples and 6 tumor samples. The procedure of picking the subsample of genes and training data is repeated 200 times, and in each run we evaluate the AUC using cross validation. When using a progressively smaller fraction of the normals, the moderated tests outperform the unmoderated tests (p-value paired signed rank test). **(B)** Joint methylation after adjusting for expression for the PKLR gene in a subset of the samples. The moderated (solid line) and unmoderated (dashed line) contour ellipses are shown. The shrinkage of variance in the tumor group is mainly caused by capping the effect of the two extreme observations using winsorization. Both the moderated test and, to a lesser extend, the unmoderated test report significant differential methylation (p-value unmoderated: 1.3 · 10^−19^; p-value moderated: 1.8 · 10^−25^). **(C)** Expression shown for two genes (MNT, SGCE) for a subset of the samples. The error bars denote the 95% interval for the moderated (solid line) and unmoderated (dashed line) estimates. The unmoderated test reports both genes as significantly differentially expressed (p-value MNT: 6.5*·*10^−4^; p-value SGCE: 4.2*·*10^−4^). However the moderated test, based on the regularized variance estimates, only reports the SGCE gene as significantly differently expressed (p-value MNT: 0.077; p-value SGCE: 3.1.10^−5^).

Winsorization can reduce the number of false positives when a location shift is driven by outliers. However, the number of false negatives can also be reduced using winsorization. For instance, if the variance is overestimated due to outliers, a true shift in location can be obscured. Figure 5B shows an example of the latter, where winsorization reduces the variance estimate of the tumor group. Albeit, testing with and without winsorization both yield a significant result, it is markedly more pronounced after winsorization, which then ultimately leads to a higher priority assignment. If the variance, on the other hand, is underestimated, small effect sizes may be rendered significant. Moderated variance estimates primarily reduce the number of false positives by shrinking the raw estimate towards the prior. The word “shrinking” can be a source of confusion; a shrunken estimate is not necessarily smaller than the regular estimate, rather it is closer to the mean of the prior distribution. Tests based on moderated variances put larger emphasis on effect size at the expense of small variance. To illustrate this, we highlight the analyses of the two genes SGCE and MNT (Figure 5C): Both genes are significantly differentially expressed using unmoderated tests, although SGCE has a markedly higher effect size than MNT. After moderating the test, only SGCE shows significant differential expression.

## 9 Conclusions and Discussion

We have presented the novel method, `EBADIMEX`, for joint identification of differential expression and methylation, and for sample classification. A major contribution of this work is the generalization of previously published moderated tests used with empirical Bayes priors for expression analysis: Here, we have developed a moderated Welch t-test for equality of means with unequal variances, as well as a moderated F-test of homogeneity in expression variance to accommodate the often highly heterogeneous nature of cancer data. Finally, we accounted for correlations between measurements and developed a moderated multivariate F-test for equality of means with equal variance, thereby extending a previously published moderated t-test for differential expression to a multivariate setting (Ritchie *et al.*, 2015). Altogether, this allows us to generalize existing empirical Bayes methods developed (Smyth *et al.*, 2004; Love *et al.*, 2014; Ritchie *et al.*,2015) for differential expression studies to a multivariate setting with multiple gene-wise measurements.

`EBADIMEX` has been applied to a TCGA breast cancer cohort of 84 normal samples and 781 tumor samples with available measures of gene expression and methylation, demonstrating the applicability of the method and its implementation. All tests applied in `EBADIMEX` are parametric, allowing for fast evaluation compared to, for instance, permutation based tests. Through label permutations and simulations, we show that the moderated tests are well behaved with uniform p-value distributions under the null.

We have demonstrated that the use of moderated tests and estimates increase statistical power, particularly for studies with few replicates or samples. In particular, we have shown that sample classification into tumor and normal benefits from making use of moderated tests and robust statistics.

The fact that methylation and expression are correlated prevents one from performing two separate analyses and combining them using Fisher’s method. Although methylation and expression measurements do not provide independent evidence of differential regulation, the use of both data types simultaneously increases our statistical power over any of the data types in isolation. This can especially prove valuable in situations where additional samples are not readily obtainable.

In the developed statistical setup, all three measurements (expression and the two methylation measurements) could be modelled by a single multivariate normal distribution. But the prior distribution of the expression variance depends on the expression level. To allow separate prior distributions for the variances of expression and methylation, we model first expression and then regress methylation on expression.

Here, `EBADIMEX` is demonstrated on cancer data, but it can just as well be applied to the other settings where we want to compare two groups to test for differential expression and methylation. More generally, the moderated Welch t-test can e.g. be used in situations where one condition shows more variability, but it is unclear which units actually have different location. The moderated F-test for equality of means can be used to include additional data types, e.g. histone modifications or chromatin state. Although `EBADIMEX` is developed with the analysis of expression and methylation data in mind, the statistical methods we develop can be applied more broadly in contexts with multiple interrelated comparisons of uni- and multivariate data.

## Author contributions

TM: Developed the statistical framework of `EBADIMEX`. Performed formal analyses and visualizations. Wrote original manuscript draft. Reviewed and edited the manuscript. MS: Data curation. Performed tests of `EBADIMEX`. MJ: Reviewed and edited the manuscript and the visualizations. JSP: Proposed the overall research idea and gave feedback on implementation and results. Reviewed and edited the manuscript and the visualizations.

## Appendix A Statistical notes

### A.1 Huber Loss

To limit the influence of a single data type (expression or methylation), when performing classification, we use the Huber Loss instead of the log-density.

If *X ∼ N* (*µ, σ*^2^), the log-density function is quadratic in *x – µ*,

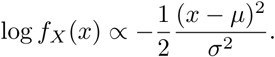

The Huber-loss function *L*(*x*) is linear when *x* is more than *k* standard deviations away from *µ*.

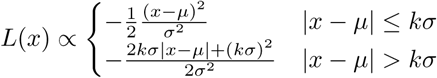

We use *k* = 2.5 in our data analyses.

### A.2 Normalization

Let *R*_*gi*_ denote the raw read count for gene *g* in sample *i*. The read counts are scaled by a sample-specific factor and then log-transformed,

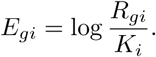

We implement two normalization strategies, namely total count (TC) and upper-quartile (UQ). In TC normalization we normalize with the total library size of the specific sample, i.e.

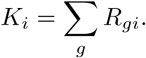

In UQ normalization we set *K*_*i*_ equal to the upper-quartile of the set *{R*_*gi*_*}*_*g*_.

The recommended method is UQ normalization (Bullard *et al.*, 2010).

### A.3 Filtering

We filter out lowly-expressed genes using a method outlined in Ding *et al.* (2015). For all genes we compute a specified quantile (by default the median, *q* = 0.5). Typically, a large group of genes display low median expression (Supplementary Figure S5). Based on a histogram we could dichotomize into two groups by manually setting a threshold. The selection of this threshold can be aided by fitting a two component normal mixture to the data: The genes having a posterior probability of belonging to the lowly expressed class larger than a half are not considered in subsequent analysis.

### A.4 Kullback-Leibler Divergence

Kullback-Leibler divergence can be used as an alternative ranking criterion to p-values. The Kullback-Leibler divergence between two normal distributions 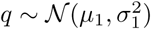 and 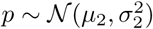 is given by

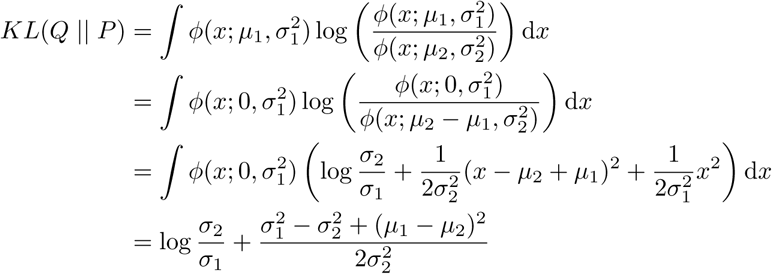

More generally, for a multivariate normal, we have

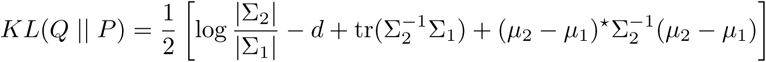

## Appendix B R-package

All functionality of `EBADIMEX` is implemented as an R-package. The package, with accompanying tutorial, is available at https://github.com/TobiasMadsen/ `EBADIMEX`.

## Supplementary figures

**Figure S1:**
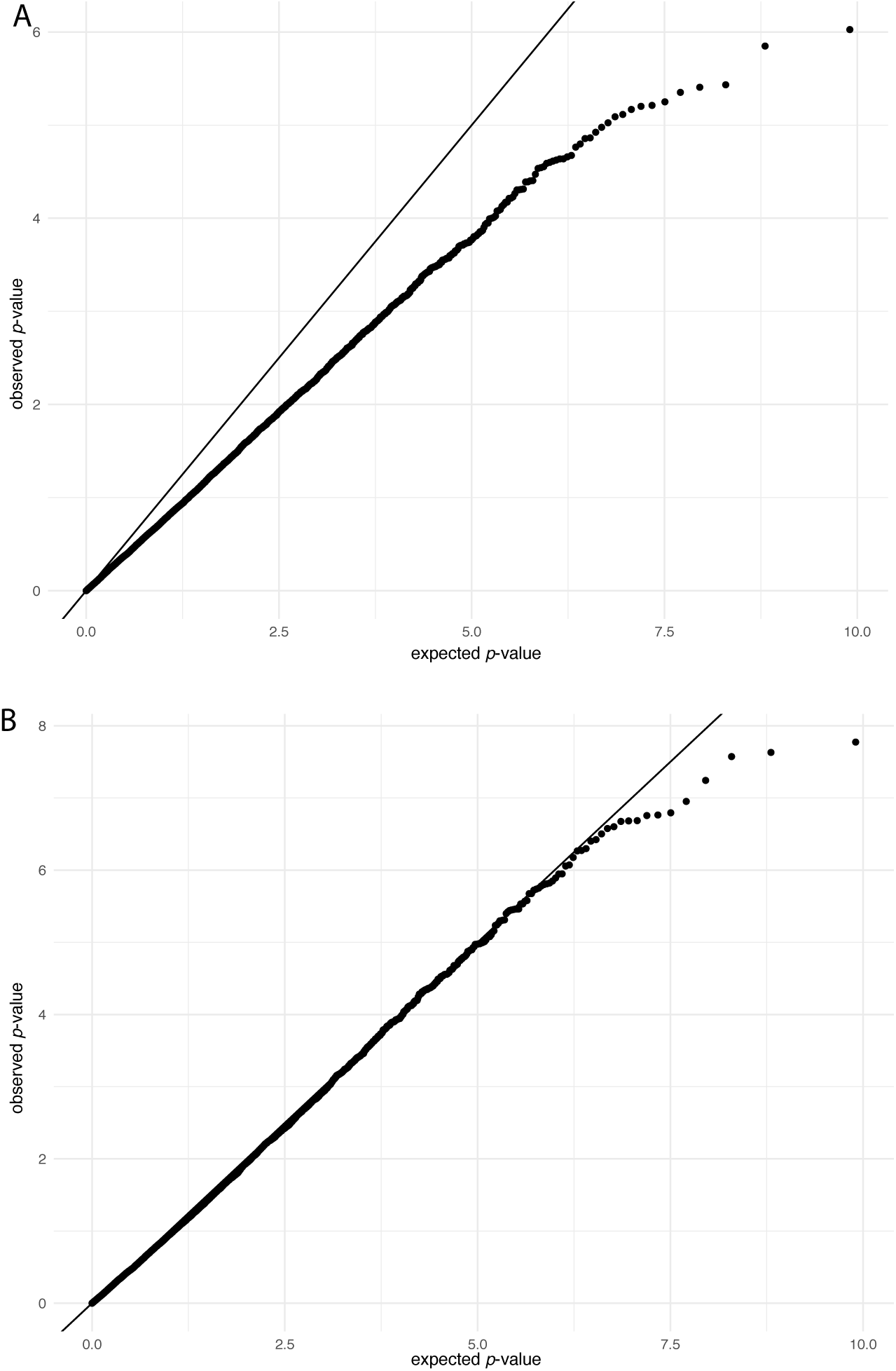
A simulation where we draw two independent variances, 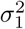 and 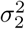 from an inverse chi-square distribution with 10 degrees of freedom. Four and six samples, respectively, are drawn from a normal distribution with mean 0 and variance 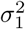 and 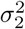. We then use the moderated welch t-test from Demissie *et al.* (2008) (panel A) and the moderated welch t-test from EBADIMEX (panel B). It is seen that the proposed t-test by EBADIMEX is nicely approximated by the t distribution.

**Figure S2:**
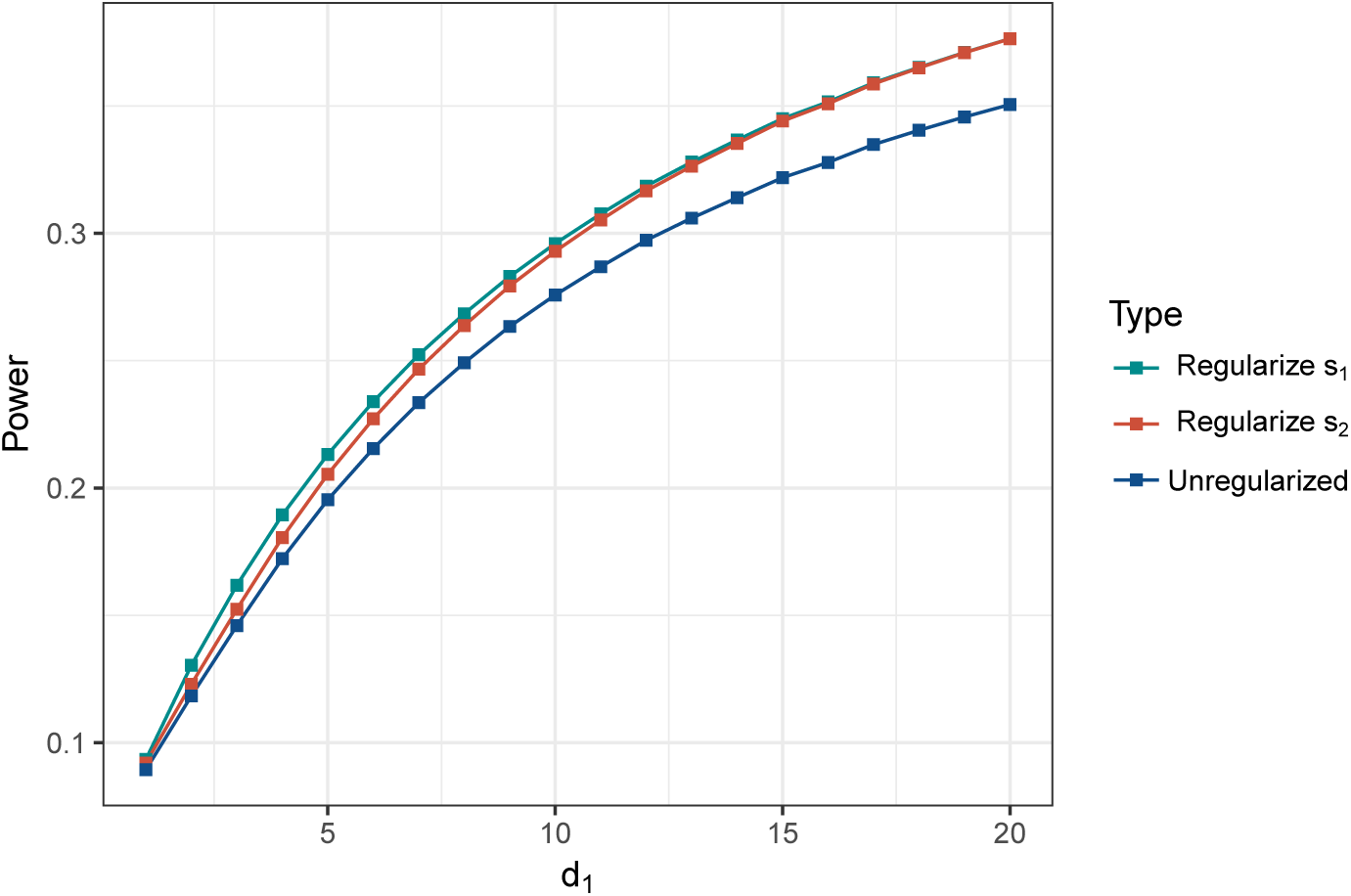
Illustration of the statistical power to reject the null hypothesis of equal variance. The two true variances are drawn from the same inverse chisquare distribution with 10 degrees of freedom, and the two sample variances 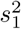 and 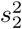 have *d*_1_ and *d*_2_ = 20 degrees of freedom, respectively. The probability of rejecting the null hypothesis is evaluated at significance level *α* = 0.05. For each datapoint, we calculated the power by simulating 10,000,000 tests. The power of the test is invariant to the choice of 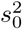 In blue is shown the unmoderated (regular) F-test, and in red and green is the moderated F test where we have moderated 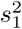 and 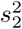, respectively. It can be seen that moderating the sample variance estimate with the lowest degrees of freedom gives higher statistical power.

**Figure S3:**
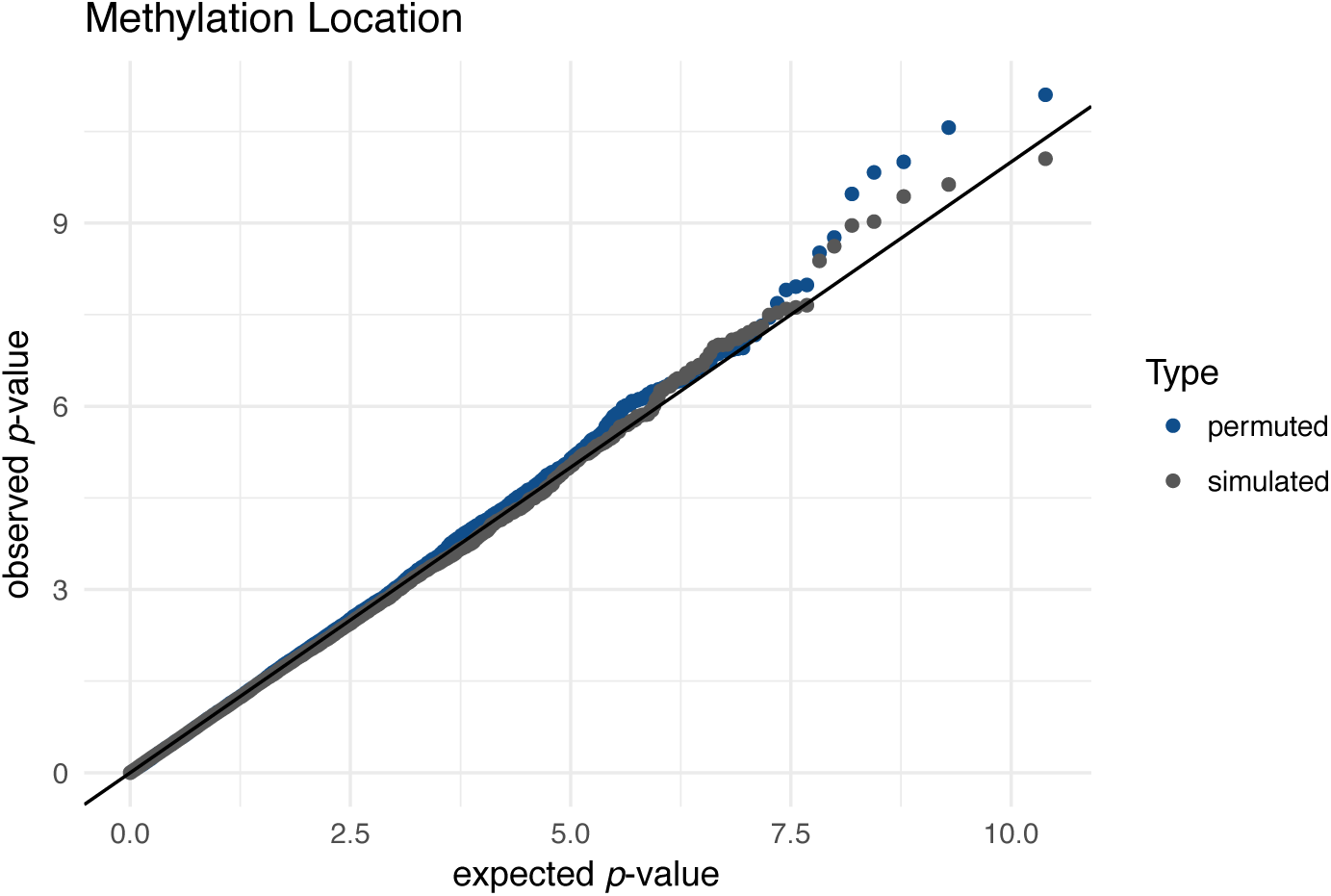
QQ-plot for the moderated F-test for location shift in the methylation data.

**Figure S4:**
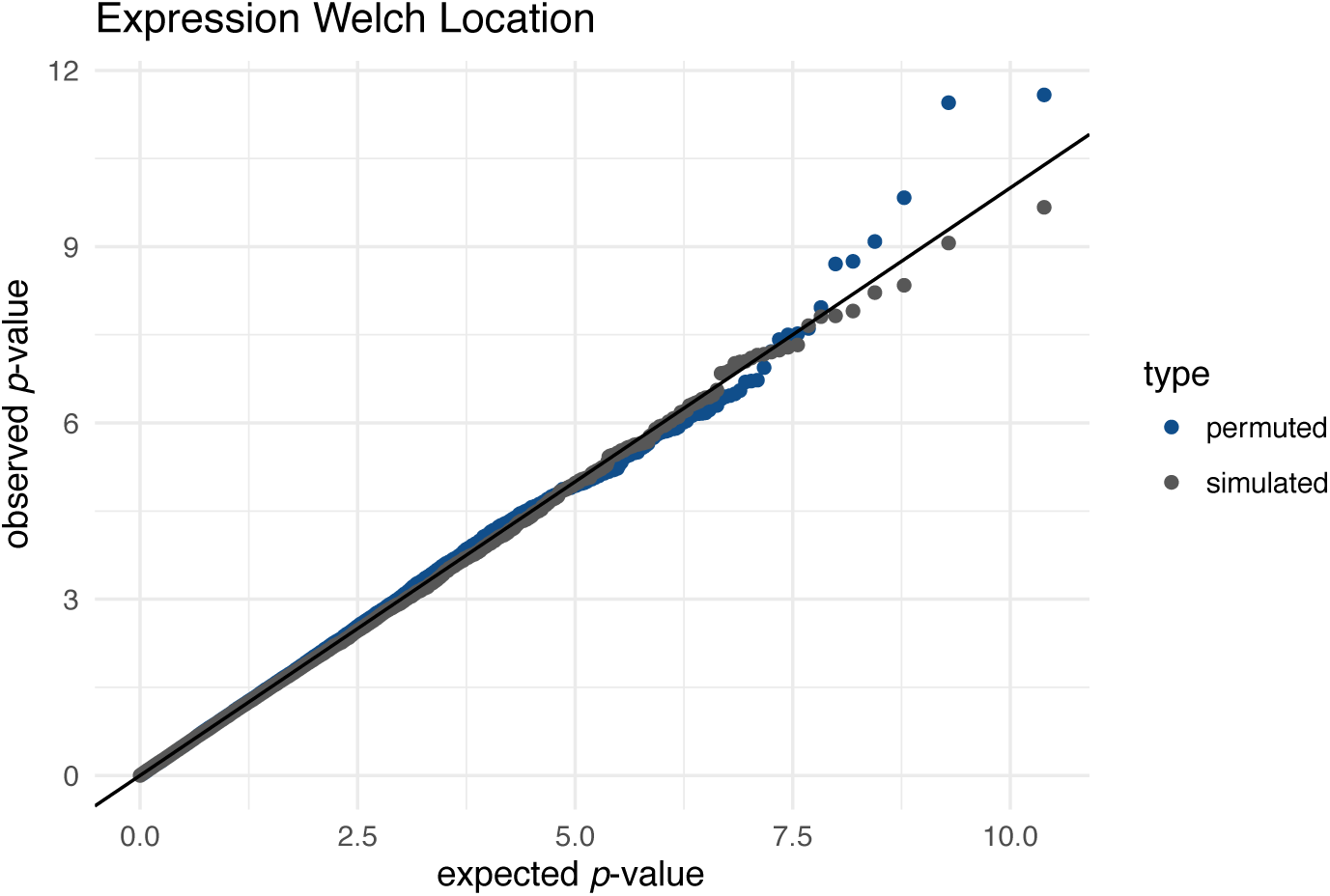
QQ-plot for the moderated Welch t-test for location shift in the expression data.

**Figure S5:**
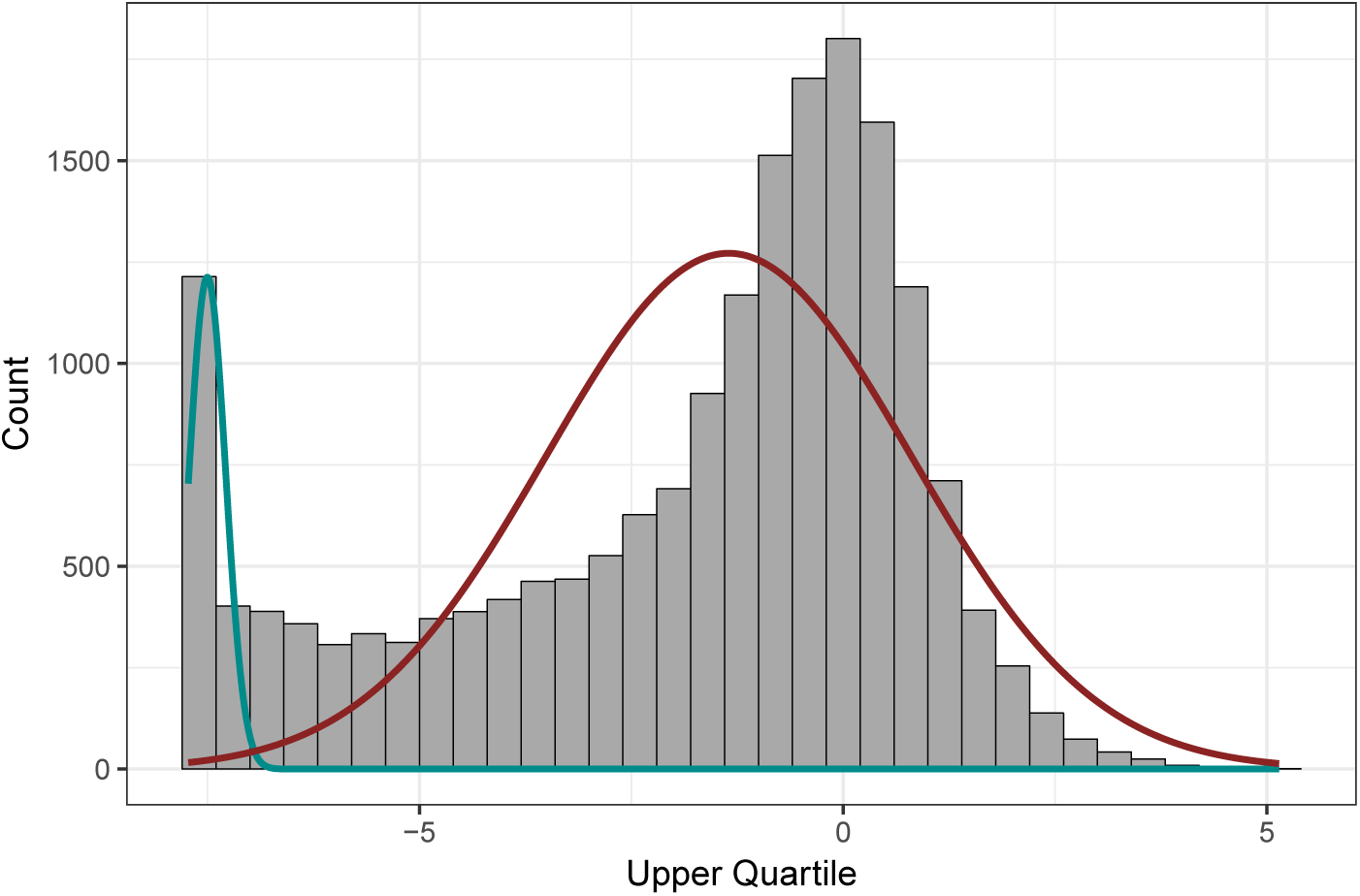
Distribution of median gene expression across all genes. A two component normal mixture is fitted to the data to identify lowly expressed genes.

